# Social Conformity Updates the Neural Representation of Facial Attractiveness

**DOI:** 10.1101/2023.02.08.527779

**Authors:** Danni Chen, Ziqing Yao, Jing Liu, Haiyan Wu, Xiaoqing Hu

**Author notes:** Corresponding Author: Dr. Xiaoqing Hu, Department of Psychology, The University of Hong Kong, Pokfulam, Hong Kong SAR, China.

## Abstract

People readily change their behavior to comply with others. However, to which extent they will internalize the social influence remains elusive. In this preregistered electroencephalogram (EEG) study, we investigated how learning from one’s in-group or out-group members about facial attractiveness would change explicit attractiveness ratings and spontaneous neural representations of facial attractiveness. Specifically, we quantified the neural representational similarities of learned faces with prototypical attractive faces during a face perception task without overt social influence and intentional evaluation. We found that participants changed their explicit attractiveness ratings to both in-group and out-group influences. Moreover, social conformity updated spontaneous neural representation of facial attractiveness, an effect particularly evident when participants learned from their in-group members and among those who perceived tighter social norms. These findings offer insights into how group affiliations and individual differences in perceived social norms modulate the internalization of social influence.

## Introduction

When observing behaviors or opinions shared by the majority, people often align their behaviors and thoughts to be consistent with others, even if initially they hold opposite views. This phenomenon, known as social conformity^1–3^, is ubiquitous: from everyday mundane choices (e.g., which movie to watch) to decisions that bear significant personal and societal consequences (e.g., whether to get vaccinated or which candidate to favor)^4–6^. Evolutionary-wise, conformity aids people in learning about uncertain environments so as to ensure survival and reproduction^7,8^. Indeed, social conformity has been documented across different species, ranging from rodents to primates^9,10^, and emerges early along the developmental trajectory^11,12^.

Despite the prevalence of social conformity, the extent to which people internalize social influence remains contentious^1,13,14^. Understanding how social influence changes one’s internal attitudes and beliefs is important: attitudes and beliefs exert powerful influences on behaviors in various settings, including consumer choices, interpersonal/intergroup relationships, and political voting, among others^15,16^. However, self-reported behaviors/opinions and internal beliefs are not always aligned, especially when external behavior could be influenced by demand characteristics and impression management strategies^17^. Thus, relying on behavioral changes to study the internalization of social influence can be challenging.

Advances are made when research leverages neuroscientific methods: If explicit behavioral changes are accompanied by changes in neural activities implicating evaluation or subjective valuation, then internalization of group influence can be inferred^2,14,18,19^. For example, during post-social influence explicit ratings on facial attractiveness, complying with social influence also enhanced neural activities in the nucleus accumbens and the orbitofrontal cortex, regions associated with subjective valuation^14^. While these studies suggested that prior social influence may induce internalization as evidenced by changed neural activity during evaluation, the explicit evaluation tasks may still be susceptible to impression management strategies. Hence, it remains unknown whether neural activity may reflect the internalization of social influence in the absence of explicit or deliberate evaluations.

In addition to investigating the internalization of social influence in the absence of explicit evaluations, it is essential to consider the source of group influence - specifically, whether people learned new information from their in-group or out-group members. People readily perceive others through the lens of social categorization, and people consistently exhibit in-group advantages in cognitive and affective processing^20,21^. From an evolutionary perspective, conforming to in-group members can be particularly adaptive, as it increases in-group homogeneity and facilitates coordination and survival^7^. Consequently, social influence from in-group or out-group members may result in varying degrees of belief updating and acceptance. However, whether people would selectively conform to in-group members remains unclear, with mixed results. Some behavioral findings suggest that people are more likely to conform to in-group norms than to out-group norms, and they may even diverge from disliked out-group members^12,22–24^. On the other hand, some studies reported no significant difference in behavioral conformity between in- vs. out-group influence or even between humans vs. computers^25–27^. Thus, how people conform to in vs. out-group influence and how group affiliations influence the associated neural activity remains an open question.

Here, we aimed to address these questions in a preregistered electroencephalogram (EEG) experiment on facial attractiveness (for preregistration, see https://osf.io/5e7kr/?view_only=cf903bc29f8543a19272046a45a8349c https://osf.io/cg6rn). In a classic social learning framework^28^, participants received normative feedback on facial attractiveness from either in-group or out-group members, introduced via a minimal group paradigm ^29–31^. We measured the changes in attractiveness ratings, which offered evidence indicative of explicit behavioral conformity. To provide evidence supporting internalization at a neural level, we devised an EEG-based face perception task in the absence of intentional evaluation or ostensible social influence.

In the EEG-based face perception task, two task features prompted us to hypothesize that we measured spontaneous evaluation of facial attractiveness. First, task-wise, the participant’s explicit task was to press a button when an object was presented on the screen, which was irrelevant to the evaluation of facial attractiveness. This task requirement thus reduced the awareness or demands of making explicit attractiveness evaluations. Second, design- and computation-wise, we developed a neural representational model that can capture spontaneous evaluations of facial attractiveness in the absence of explicit evaluation. While previous studies have found that event-related potentials (ERP), such as the face-sensitive N170 and evaluation-related late potential component (LPC) ^32–36^, can indicate perceived facial attractiveness, there were limitations. The extent to which ERPs indicate facial attractiveness remains mixed. For example, when participants made gender judgments, there were no significant differences in ERP between attractive and non-attractive faces^37^. Additionally, there are considerable individual differences in attractiveness perception, which may reduce ERP’s sensitivity in assessing one’s attractiveness perception^38–40^. To overcome these shortages, we applied multivariate neural representation similarity (RSA) analyses to the face-elicited EEG to extract neural representations of facial attractiveness^41^, which could capture complex neural representational patterns across multiple channels. Notably, our task also included prototypical attractive faces. By computing the neural representation similarities between the learned faces and the prototypical attractive faces, we could infer whether the learned faces were perceived as more or less “attractive” as a result of social influence. Importantly, this RSA approach allowed us to build a sensitive and individualized neural representation model of perceived attractiveness, even when univariate neural activity fails to show differences^42^.

We preregistered our hypotheses that participants would be more likely to behaviorally comply with in-group than out-group opinions, as evidenced by explicit attractiveness rating change (Hypothesis 1). Concerning attractiveness-related ERPs and neural representations updating, we aimed to test two competing hypotheses: Participants would only internalize in-group members’ influence (Hypothesis 2a), or they would internalize both in- and out-group influence (Hypothesis 2b) as evidenced by spontaneous neural representations of facial attractiveness. Considering the individual differences in the propensity to conform to other^43–45^, we explored how individual differences in their perceived tightness-looseness of social norms may modulate the behavioral and neural effects of social compliance^46,47^.

## Results

### Preregistered Confirmatory Behavioral Results

Forty-eight participants (37 females, 43 heterosexuals, age, *mean* = 23.98, *S.D.* = 3.13) were recruited from a local university, among which 45 participants were included in the EEG analyses. Participants visited the lab twice, separately by seven days. In the first lab visit, participants completed a series of questionnaires followed by computer-based tasks. For computer-based tasks, participants completed three phases: 1) pre-learning, 2) learning, and 3) post-learning (Figure 1). In the pre-learning phase, participants performed a face perception task and an explicit rating task as baseline measures, with 60 medium-attractive to-be-learned faces, 10 medium-attractive no-learning control faces, and 10 prototypical attractive faces. In the face perception task, participants viewed 480 faces intermixed with 144 objects, divided into 6 blocks, with brainwaves being recorded. Participants were instructed to press buttons when they saw objects, which were designed to ensure their attention was maintained throughout the experiment. In the explicit rating task, participants rated each of the 80 faces with a mouse on attractiveness, confidence, perceived competence, and perceived warmness (1 to 11). During the learning phase, participants were randomly assigned to one of two groups (Green or White) while being told the assignment was based on their shared attribution with the group members (i.e., a minimal group manipulation)^29,30^. Next, participants learned about the attractiveness ratings feedback from either in-group or out-group members (i.e., Affiliation), which was either Higher, Lower, or Consistent (i.e., Feedback) than/with their initial ratings, resulting in a 2 (Affiliation) by 3 (Feedback) within-subject design with 10 experimental faces in each condition. The sources of the feedback were indicated by the color of ticks on a 1-11 scale. After the learning task, participants performed a repeated face perception task and an explicit rating task, which was used to compute updates of facial attractiveness ratings.

**Figure 1.**
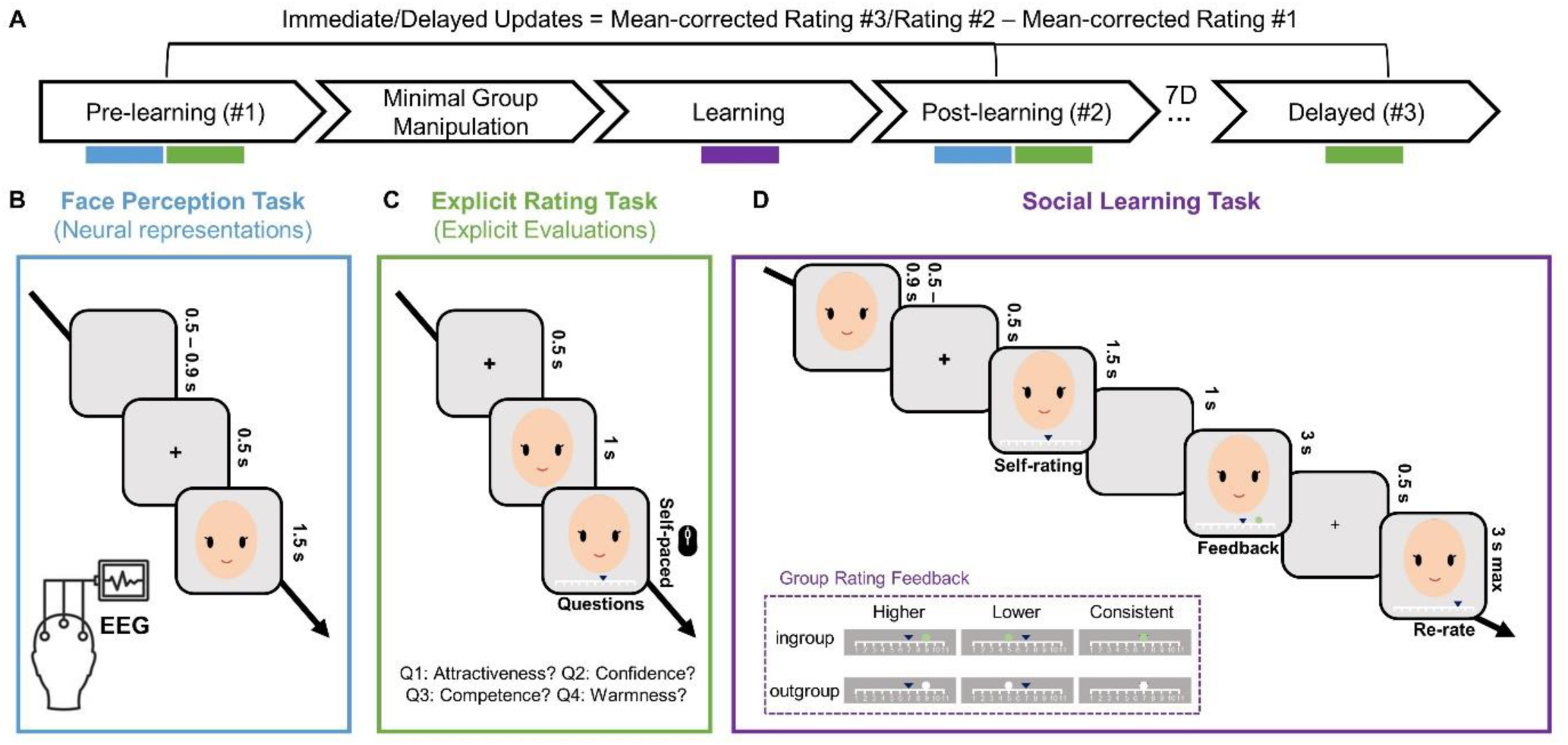
Experimental Procedure. (A) The upper row represents the procedural flow, with colored rectangles below to illustrate each task in sequence. In the pre-learning and post-learning phases, participants completed the same EEG-based face perception task (B, blue rectangle) and the behavioral explicit rating task (C, green rectangle). In the delayed phase, participants only completed the behavioral explicit rating task. Between the pre- and post-learning phases, participants completed the minimal group manipulation to learn about their group affiliations. In the social learning task, participants learned the attractiveness ratings of each face from either in-group or out-group members (D, purple rectangle). The face icon illustrated the Asian female facial stimuli used in our experiment, with hair and ears removed from the faces.

We first examined whether group affiliation interacted with social influence in updating attractiveness ratings. To account for the regression-to-mean effect and potential systematic rating differences across different phases (pre-learning, post-learning, and delayed), we calculated the mean-corrected attractiveness ratings for each participant and each face stimulus at each phase. We first calculated the average attractiveness rating of all faces within the corresponding phase, and then we subtracted this average rating from the individual ratings to obtain the mean-corrected attractiveness rating ^48,49^. We computed the attractiveness update by subtracting the pre-learning mean-corrected rating from the post-learning (immediate update) and delayed mean-corrected rating (delayed update) for each individual face.

An affiliation (in- vs. out-group) by feedback (higher, lower, consistent) repeated measures ANOVA on the immediate update of attractiveness ratings (post-minus pre-learning) revealed a significant feedback effect (*F* (1.91, 89.75) = 9.17, *p* < .001, *η*^2^ = 0.07, BF_10_ = 720.31, Figure 2A). Participants rated the faces less attractive in the lower condition than in the higher (*t* (47) = 3.79, *p* = .001, *d* = 0.60, BF_10_ = 109.59) and in the consistent condition (*t* (47) = 3.56, *p* = .003, *d* = 0.53, BF_10_ = 47.22), while there was no significant difference between the higher and consistent conditions (*t* (47) = 0.89, *p* = 1.000, *d* = 0.14, BF_10_ = 0.17). However, neither the main effect of affiliation (*F* (1, 47) = 0.31, *p* = .579, *η*^2^ = 0.001, BF_10_ = 0.15) nor the affiliation by feedback interaction was significant (*F* (1.93, 90.52) = 0.66, *p* = .513, *η*^2^ = 0.005, BF_10_ = 0.20), with Bayesian factors strongly favoring the null hypothesis. The same analysis on the delayed updates of attractiveness ratings (delayed minus pre-learning) revealed no significant effect (*p*s > .430, BF_10_s < 0.09, Data S1). Taking the regression to mean effect into consideration, we repeated the analyses using faces that were matched on baseline ratings across feedback conditions^14,48^ and obtained similar results (Data S2).

**Figure 2.**
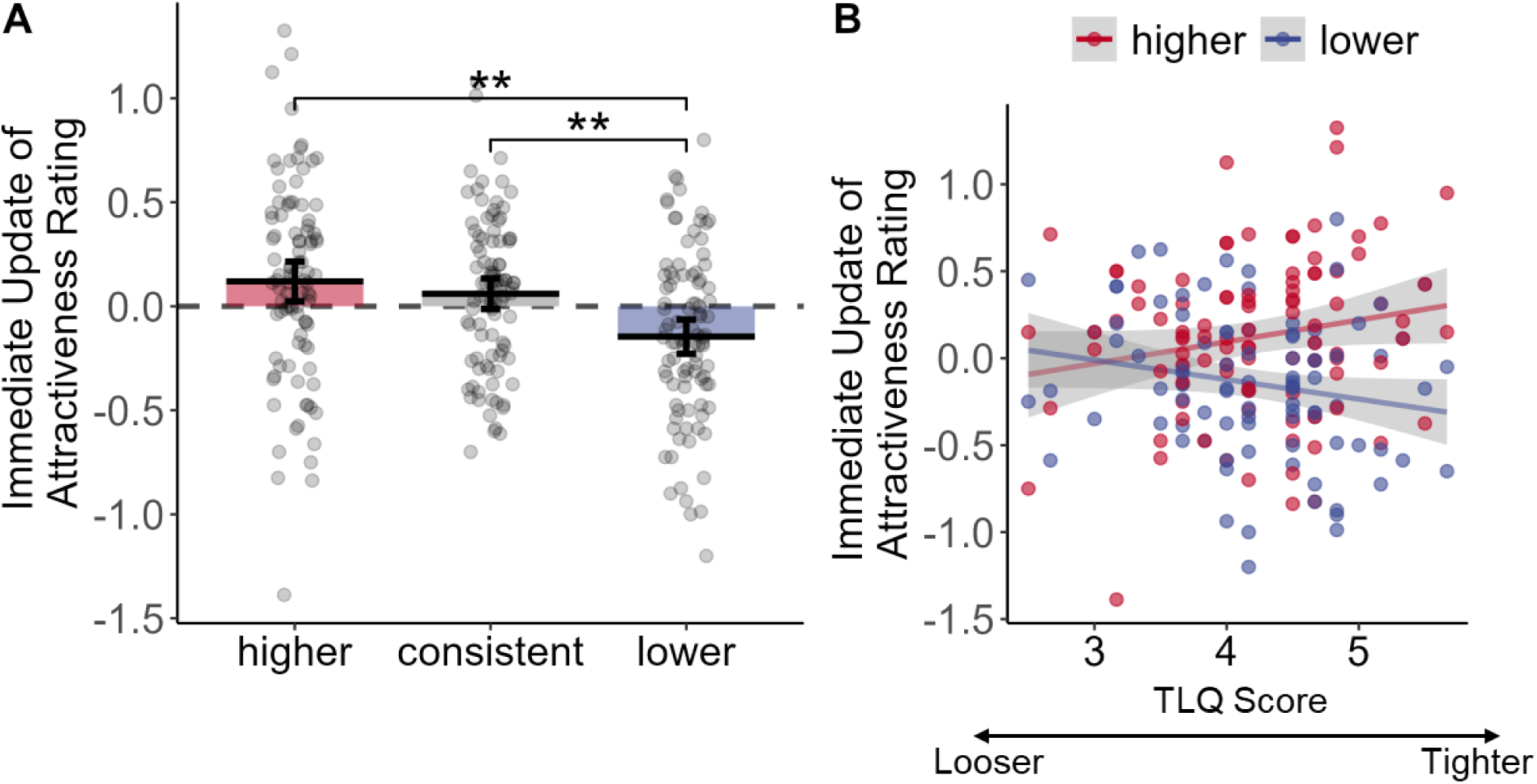
Behavioral results from preregistered analyses. A) Social influence induced the update of attractiveness ratings across in-group and out-group influences from pre- to post-learning tests. For feedback by affiliation results, see Figure S1. For the impact of continuous rating discrepancies on explicit rating changes, see Figure S2. B) Individuals who perceived tighter social norms (higher scores on the x-axis) showed stronger immediate updates in explicit ratings (i.e., higher > lower) than individuals who perceived looser social norms.

Together, these results suggested that social influences induced changes in explicit evaluations at least in the immediate test. However, in contrast to Hypothesis 1, the group affiliation did not significantly differ in the updates of explicit ratings.

### Preregistered Exploratory Behavioral Results

Next, we examined the correlations between individual difference variables (e.g., perceived tightness-looseness, empathy, socially desirable responding, and social phobia) and immediate and delayed updates of attractiveness rating, respectively (for results, see Table S2). Among these individual difference measurements, we observed that perceived tightness-looseness (TLQ score) showed opposite directions in predicting immediate updates for higher and lower feedback conditions, respectively. We thus conducted a moderation analysis using TLQ, feedback (higher vs. lower), and their interaction as independent variables, and the immediate updates in attractiveness rating as the dependent variable. The regression model showed a significant interaction between the TLQ scores and feedback conditions on the immediate updates of attractiveness ratings (*b* = −0.24, SE = 0.09, *p* = .009; Figure 2B). The post-hoc simple slope analyses revealed that the standardized regression coefficient for the higher condition was significantly higher than for the lower condition: *t* (188) = 2.65, *p* = .009. Particularly, the standardized regression coefficient in the higher and lower conditions showed opposite effects (higher condition: *t* (188) = 1.97, *p* = .050; lower condition: *t* (188) = −1.77, *p* = .078). The significant interaction suggests that individual differences in perceiving the social norms significantly modulated the explicit compliance effect, per feedback directions.

### Preregistered Confirmatory ERP Results

In the face perception task, we examined pre- vs. post-learning changes of the face-sensitive N170 on the pre-defined left and right occipitotemporal sites, and of the evaluation-related LPC on pre-defined central-parietal and frontal-central sites (for ERPs, see Figure 3). The affiliation by feedback repeated measures ANOVAs showed no significant effects of affiliation (*p*s > .056, *η*^2^ < 0.008), of feedback (*p*s > .176, *η*^2^ < 0.005), or their interaction (*p*s > .088, *η*^2^ < 0.009, full results are provided in Table S3). Thus, the effect of social influence on facial attractiveness did not emerge when using univariate ERP analyses in N170 and LPC.

**Figure 3.**
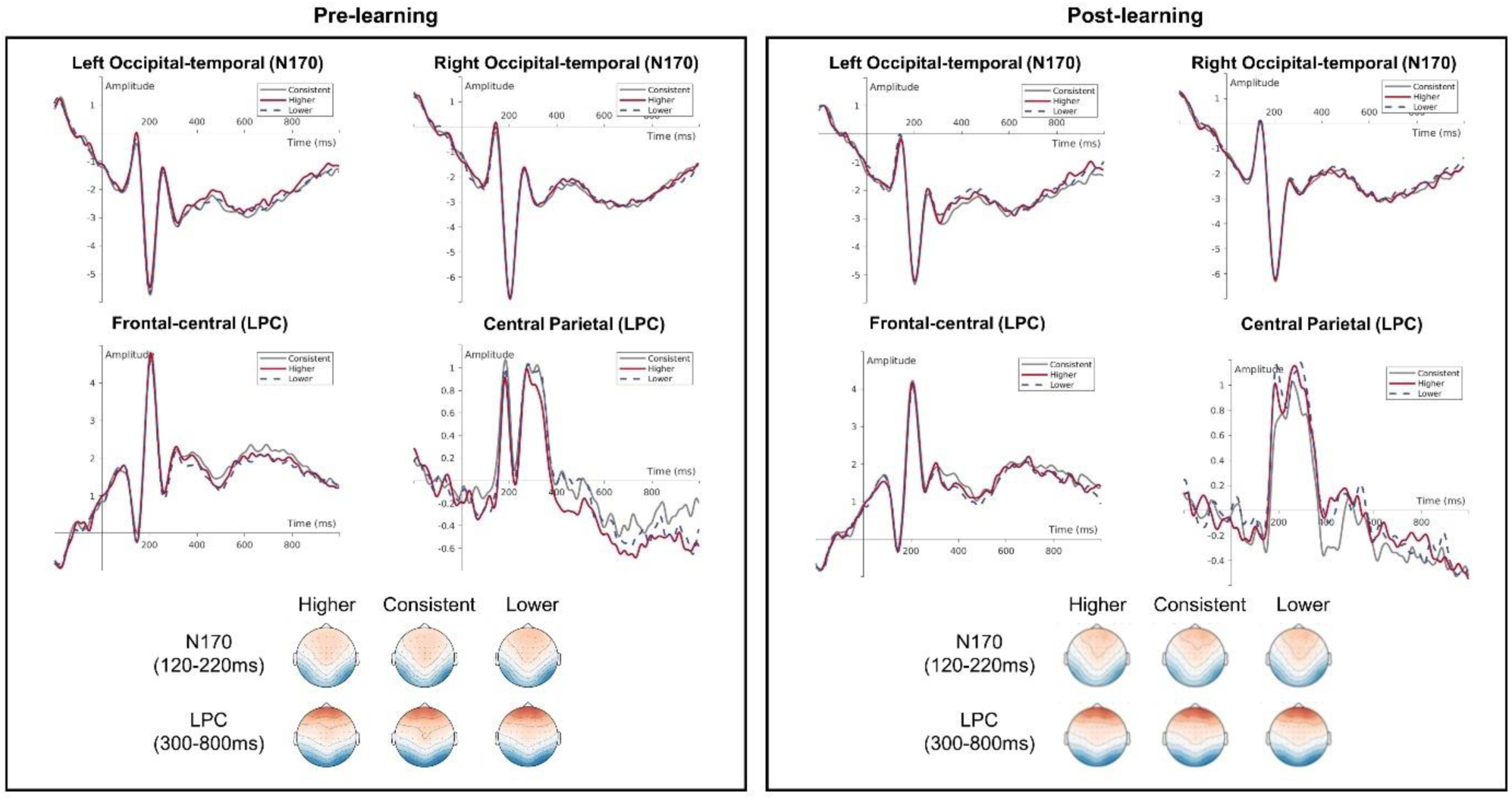
Face-locked ERPs and topography in the pre-and post-learning phases. CP: central-parietal sites (CPz, CP1/2, Pz, P1/2); OT: occipitotemporal sites (left: T7, TP7, P7, PO7; right: T8, TP8, P8, PO8); FC: frontocentral sites (Fz, FCz, F1/2, FC1/2). We examined N170 at the left and right occipital-temporal sites, and LPC at the frontal-central and central-parietal sites.

### Preregistered Exploratory RSA Results

Given the limitation of univariate analysis in analyzing multidimensional information^42^, we further examined the spontaneous neural representations of facial attractiveness using multivariate RSA. Here, we calculated the multivariate neural representation similarities between the experimental face and prototypical attractive faces, i.e., the experimental-prototype face similarity (EPS) as an individualized neural index of attractiveness (Figure 4A).

**Figure 4.**
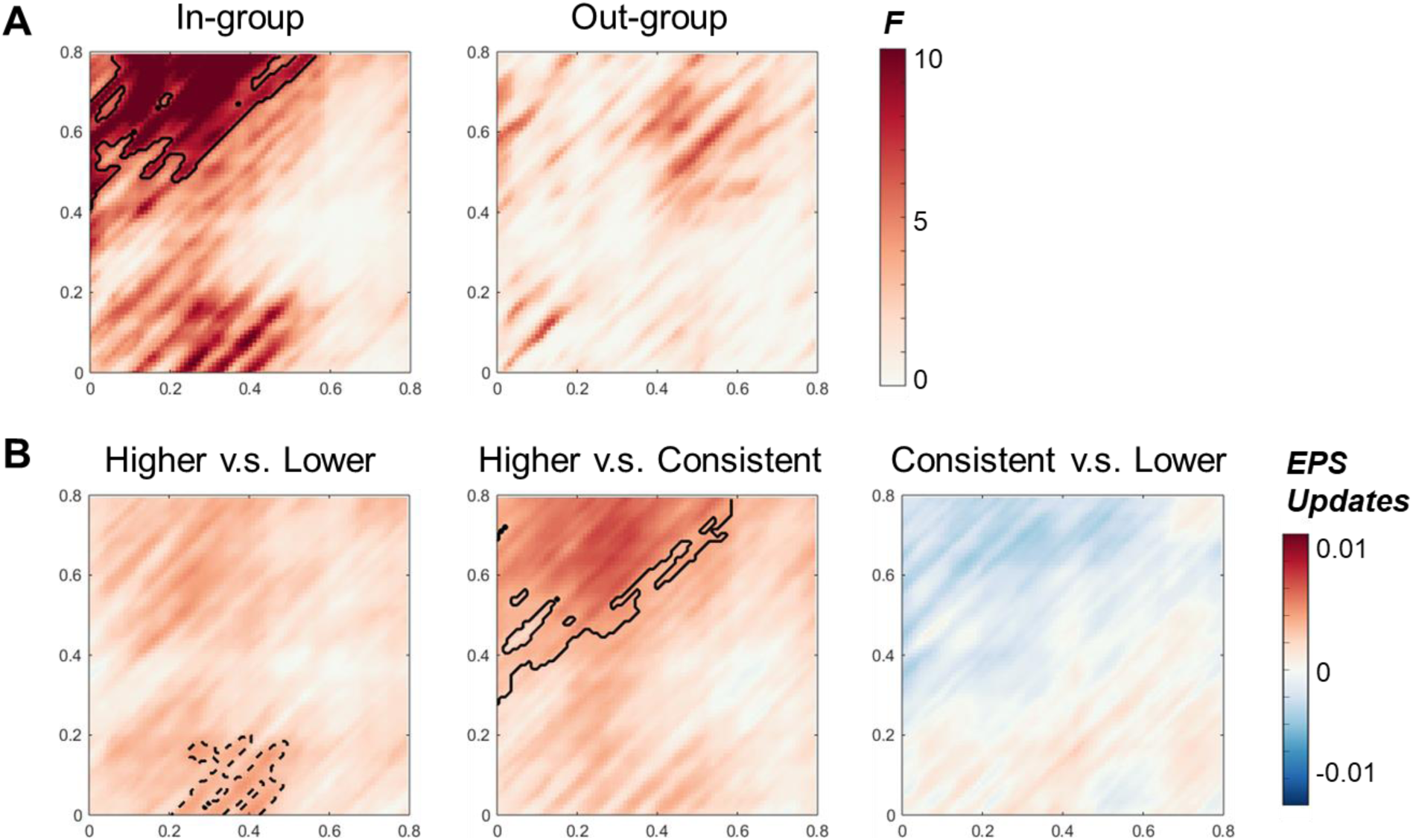
RSA Results. A) The F-value of the one-way repeated-measure ANOVA of feedback on the pre- to post-learning EPS updates, with cluster-based permutation analysis testing on EPS update for in- and out-group conditions separately. A significant feedback effect in EPS update only emerged in the in-group condition. B) EPS updates between different feedback contrasts in the in-group condition using paired t-tests with cluster-based permutation tests, in which we observed a significant cluster in the higher vs. consistent contrast. The X-axis shows the timescale of the learned experimental faces, and the Y-axis shows the timescale of the prototypical beauty faces. The solid contour indicates significant clusters (*p*_cluster_ < .05). The dashed contour indicates clusters with .05<*p*_cluster_ < .10.

To better control the pre-learning baseline EPS, we examined the effect of feedback on the EPS update from pre- to post-learning (i.e., EPS of post-learning minus pre-learning phases) across in- and out-group conditions. The results showed that feedback significantly impacted the EPS update (*p*_cluster_ = .013, cluster-based permutation test; for details, see Methods section). Post-hoc analysis on the EPS update revealed that the higher feedback condition was associated with a significantly higher EPS update than the consistent condition (*p*_cluster_ = .005). No significant higher vs. lower difference, or consistent vs. lower difference, was found (*p*_cluster_s > .197).

We next examine the effect of feedback on the EPS update from pre- to post-learning for in-group and out-group conditions, respectively. In the in-group condition, feedback significantly modulated the EPS update (*p*_cluster_ = .048; Figure 4A left). Post-hoc analysis on the EPS update showed that the higher feedback condition was associated with significantly higher EPS update than the consistent condition (*p*_cluster_ = .016; Figure 4B center) and numerically higher EPS update than the lower condition (*p*_cluster_ = .070; Figure 4B left). No significant difference between the consistent and the lower conditions on the EPS update was found (*p*_cluster_s > .260; Figure 4B right). In contrast, no significant effect of feedback on the EPS update was found in the out-group condition (*p*_cluster_s > .211, Figure 4A right). When conducting an affiliation by feedback repeated measure ANOVA on the EPS update, we did not find a significant affiliation by feedback interaction (*p*_cluster_s > .530). Additionally, when examining EPS in the pre- and post-learning phases separately, we did not find significant effects of feedback and affiliation on the EPS (*p*_cluster_s > .067; Data S3). Further exploratory analyses showed that among female participants, the EPS updates from the higher vs. lower feedback contrast were significantly higher in the in-group than in the out-group condition (*p*_cluster_s < .041, Figure S3).

Having shown that the perceived tightness-looseness influenced the attractiveness rating change at the behavioral level, we further explored whether the perceived tightness-looseness would influence the spontaneous neural representations of facial attractiveness. To this end, we divided participants into high (*n* = 22) and low (*n* = 23) tightness-looseness groups (TLQ) based on whether their perceived tightness-looseness was higher than the median tightness-looseness score. As we found that the EPS updates were significant in the in-group condition, we focused on EPS updates of the two sub-groups in the in-group condition. The cluster-based permutation test found that for the high-TLQ sub-group, the pre- vs. post-learning EPS update (Figure 5) of the higher condition was significantly higher than that of the lower condition (*p*_cluster_ = .028) and of the consistent condition (*p*_cluster_ = .028). No significant cluster was found in the consistent vs. lower contrast. In contrary, for the low-TLQ sub-group, no significant cluster was found among all contrasts (*p*_cluster_s > .101). We confirmed that no significant difference between the high- and low-TLQ group in the in-group favoritism ratings (Data S4). Again, this effect was particularly evident in the in-group condition (see Figure S4 for results in the out-group condition). These results suggested that the impact of social influence on spontaneous neural representation of facial attractiveness was likely driven by participants who perceived tighter social norms.

**Figure 5.**
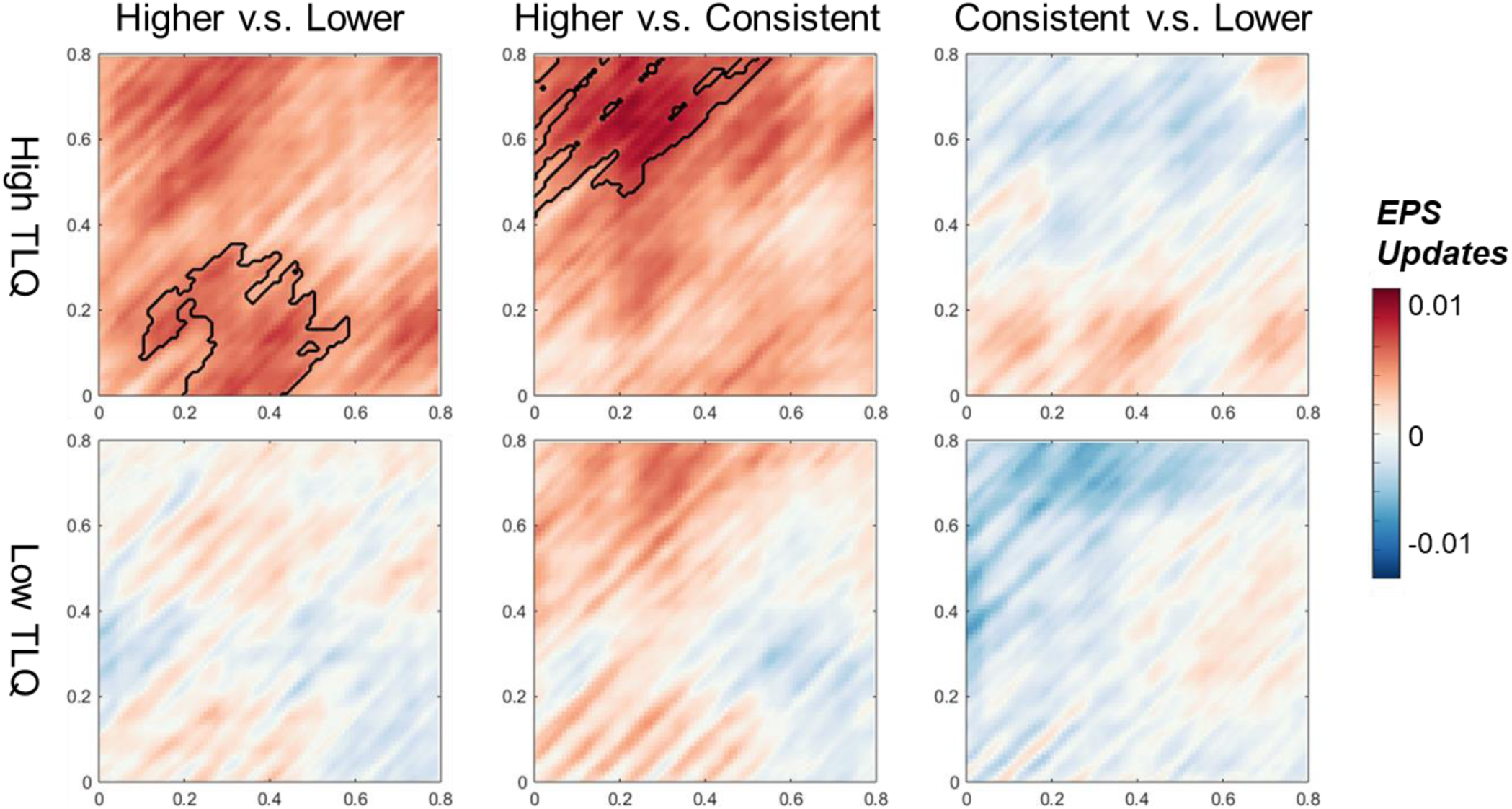
EPS update in the in-group condition of the high and low tightness-looseness (TLQ) sub-groups. The upper row is the EPS update in the High TLQ sub-group, in which we found significant clusters in the higher vs. lower, and the higher vs. consistent contrast, but not in the consistent vs. lower contrast. The lower row is the EPS update in the Low TLQ sub-group in which no significant clusters were found. The solid contour indicates the significant clusters (*p* < .05).

## Discussion

Evolutionary-wises, following the crowd bears significant survival benefits. Employing a social learning paradigm, our preregistered EEG study examined social conformity after people learned from in-group and out-group consensus on facial attractiveness. Behaviorally, we found that both in- and out-group influence changed explicit attractiveness ratings. To quantify the internalization of social influence on face attractiveness evaluation, we leveraged the computation power of representational similarity analysis (RSA) to extract neural representations of facial attractiveness when social influence is no longer salient. We showed that social conformity updated the spontaneous neural representations of facial attractiveness, suggesting internalization. Notably, neural representational update was particularly evident when participants learned from their in-group members, and among those who perceived tighter social norms.

Regarding the explicit behavioral changes, we replicated previous findings that people would change explicit attractiveness ratings to comply with others^14,48,50^. Moreover, participants changed their attractive ratings in response to both in- and out-group influence, and we did not find an in-group advantage effect on explicit ratings^12,22–24^ per our preregistered Hypothesis 1. The lack of an in-group advantage effect was also supported by the Bayesian Factors that strongly favor the null over the alternative hypothesis. Differences in the group affiliation manipulation between the current and previous experiments might explain the lack of in-group advantage observed here. In our study, we adopted a minimal group paradigm in which participants were randomly assigned to arbitrary groups (White or Green), while previous research used real-life group identities (e.g., Chinese vs. American^23^; Caltech students vs. Sex offenders^12^) to highlight group membership. Using real-life group identities resulted in higher compliance to in-group than to out-group members, i.e., an in-group advantage effect, probably due to their higher motivational salience than the group affiliations formed in lab-based minimal group paradigms. Indeed, using a similar minimal group manipulation, a previous study also showed that people conform to both in- and out-group members, and an in-group advantage effect only emerged when oxytocin was given^25^. These findings raised the possibility that the evaluation updating in our experiment could be a result of any learning or even due to re-evaluation. Given that we did not include a non-social control condition, we could not fully address the question of social vs. non-social learning. However, our results still favor a social learning account: evaluation updating was sensitive to participants’ perceived tight-looseness of social norms, which highlights the social nature of evaluation updating. To gain a deeper understanding of the mechanisms underlying social conformity, future research shall include a non-social feedback group (e.g., human vs. computer ^27^) to better distinguish the social vs. non-social impact on evaluation updating.

The investigation of the spontaneous neural representations of facial attractiveness during face perception may provide insight into whether participants internalize social influences. Specifically, by applying RSA to each participant’s EEG elicited by prototypical attractive faces, we were able to build an individualized neural representation model of prototypical attractiveness, which would be compared with the experimental faces. The analytical power of RSA, combined with the face perception task, allows us to detect subtle changes in the neural representation of facial attractiveness, thus capturing the internalization of social influence even in the absence of ostensible social influence and explicit attractiveness judgments. Our findings suggested that social conformity updated the spontaneous neural representation of facial attractiveness. Moreover, these updated neural representations happened at an early time window, overlapping with the time windows during which face perception^33,37,51^ and attractive evaluation often occur ^41^. Together, our results provide a mechanistic explanation for continued social influence: If social influence changes the spontaneous neural representation of stimuli, the influence can change behavior even when social influence or norms are not salient. Lastly, although updating of spontaneous representation was significant after people learned from their in-group members, the critical group by feedback interaction was not significant. This prevents us from concluding that participants would preferentially internalize social influence from their in-group members. Future research shall examine the updating of neural representations using real-life group identities that bear higher ecological validity (e.g., one’s nationality^23^).

People differ in how they perceive social norms^47,52^. One dimension of norm perception is the perceived tightness-looseness of social norms, which represents how an individual perceives society as having tight or loose norms and having low or high tolerances for norm-deviant behaviors^46^. Even within the same cultural context, perceived tightness-looseness would vary across individuals and would impact how they react to social influence^47^. For the explicit attractiveness ratings, we found that those who perceived tighter social norms were associated with increased levels of explicit attractiveness rating changes. Intriguingly, perceived tightness-looseness norms also modulated attractiveness perceptions at a neural level: those who perceived tighter norms showed stronger changes in the spontaneous neural representations of facial attractiveness. Recent research also found that perceived tightness-looseness of social norms predicted the amplitude of N400, an ERP component sensitive to semantic incongruity, when participants viewed various norm-deviant behavior^52^. Extending this research, our results suggested that those who perceived tighter social norms would be more intrinsically motivated to follow social influence and showed higher levels of internalization, as evidenced by both behavioral and neural representation changes towards in-group influence.

Our study demonstrated that learning from in- and out-group members modulated attractiveness perception. In a broader sense, social perception can be modulated via multiple processes tapping into social-motivational-affective mechanisms. For instance, the likability of neutral faces could be reduced when they were paired with unrelated negative information, even when such affective stimuli were presented unconsciously^53,54^. Our research joins this effort, contributing to our understanding of how to modulate social perceptions including perceived attractiveness, likability, and trustworthiness^28,55^. Future research can employ the task and analytical approaches (e.g., prototypical faces in the face perception task, the RSA) to investigate how social/affective manipulations can alter social perception at a neural representation level^42,56^.

Limitations and future directions shall be discussed. First, we did not record EEG during the delayed test, which restricted us from investigating the longevity of internalization at the neural level. As attitude and behavior are not always aligned, future research should focus on the long-term effect of neural representational changes. Second, our research focused on facial attractiveness, which might be easier to challenge than topics that are central to one’s values and worldview, such as moral values and political views. Future research could apply this social learning framework, combined with the neural representation approach, to examine how social influence would change attitudes and beliefs that are core to one’s worldviews and values. Third, it is important to note that there could be gender differences in facial attractiveness perception, social conformity, and intergroup biases at both the behavioral and neural levels ^57–59^. Given that most of our participants were heterosexual females and we only included Asian female faces, this may limit the generalizability of our findings. Investigating potential gender differences using similar paradigms is warranted in future studies. Finally, while we measured individual differences in perceived tightness-looseness among participants from the same culture, future research shall consider cross-cultural studies to examine how tight-looseness culture may influence explicit and spontaneous conformity.

To conclude, our preregistered EEG study found that social compliance and internalization can happen even without overt normative feedback and intentional evaluation. Notably, this effect was particularly evident when people learned from their in-group members, and among those who perceive tighter social norms. Given that social compliance has survival benefits, future research shall further investigate how conformity and internalization of social influence may build up norms abided by group members.

## Methods

### Participants

We preregistered to recruit 42 participants, which is larger than previous similar EEG experiments on attractiveness^32^ and social conformity^60–62^, and allows us to detect effect sizes in the range of 0.40 to 0.50^1^. Anticipating potential attrition and data exclusion, we recruited 48 participants (37 females; 43 heterosexuals; age, mean = 23.98, S.D. = 3.13) from a local university. Participants received monetary compensation at a rate of 80 HKD/hour. Three participants were excluded from subsequent EEG analysis due to excessive EEG artifacts, resulting in 45 participants who met our preregistered inclusion criteria: 1) Following artifact rejection, each participant’s clean EEG segments should be more than 50% of total trials in the face perception task in both pre- and post-learning phases; and 2) participants should correctly report their assigned group identity. All participants were native Chinese speakers, right-handed, not color blind, had a normal or corrected-to-normal vision, and did not report any history of neurological or psychological disorders. All participants provided written informed consent prior to the participation and were debriefed and compensated after completing the study. This research procedure was approved by the Human Research Ethics Committee of the University of Hong Kong (HREC No. EA1912003).

### Materials

We selected 113 photographs of East Asian female faces from previous research ^64^. Additionally, we generated 21 morphed faces by morphing four randomly selected faces from the same face database by FunMorph. The morphed faces would serve as the prototypical attractive faces because people perceive faces as more attractive when they are closer to the prototype ^65,66^. For faces, hair and ears were manually removed from the faces by Adobe PhotoShop. All photos were round-cropped and manually aligned with size, luminance, lightness, and color using Adobe Lightroom.

We conducted a pilot study to select medium-attractive face stimuli. An independent group of participants (*n* = 18, college students) rated the attractiveness of each of the 134 faces (including both morphed and original faces) on a 1-7 scale. Next, we selected 70 experimental faces and 10 prototypical beauty faces based on their average attractiveness ratings.

Specifically, for the 113 original faces, we first removed 29 faces with averaged attractiveness ratings greater or less than 1.5 standard deviations (S.D.). Within the remaining face stimuli, we removed 14 faces with the highest attractiveness ratings so that we could retain 70 medium-attractive East Asian Female faces for 10 faces in each experimental condition. We removed the top-rated attractive faces so that the to-be-learned faces could be more distinct from the morphed faces in terms of attractiveness. For the 21 morphed faces, we selected 10 faces with the highest attractiveness ratings to serve as prototypical attractive faces.

Together, 70 medium-attractive and 10 highly attractive morphed East Asian female faces were retained in the formal experiments. Data from the pilot participants confirmed that the prototypical faces were significantly more attractive than the median-attractive faces (of the 7-point scale; prototype faces, *mean* = 5.84, *S.D.* = 0.66; target faces, *mean* = 3.77, *S.D.* = 0.69; *t* (17) = 11.91, *p* < .001, *d* = 3.09; details see Data S5).

We only included medium attractive Asian female faces in the current experiment because 1) the medium attractive faces provided participants with greater flexibility to adjust their ratings (e.g., increase or decrease) in the social learning task, and 2) the inclusion of only the female faces can control the potential gender differences in face perception. This approach is in accordance with previous studies with similar designs^14,48,50^.

### Procedure

All tasks were programmed and presented by PsychoPy (version 2020.1.3)^67^. Participants visited the lab twice, separated by seven days. In the first lab visit, participants completed the Positive and Negative Affect Schedule (PANA-SF)^68^, Interpersonal Reactivity Index (IRI)^69^, Tightness-Looseness Questionnaire (TLQ)^46^, Socially Desirable Responding (SDR)^70^, and Social Phobia Inventory (SPIN)^71^, followed by computer-based tasks.

For computer-based tasks, participants completed three phases: 1) pre-learning, 2) learning, and 3) post-learning (Figure 1). Participants performed a face perception task and an explicit rating task in both the pre-learning and post-learning phases, with 70 medium-attractive experimental faces intermixed with 10 prototypical attractive faces. In the face perception task, participants viewed 480 faces intermixed with 144 objects divided into 6 blocks (for trial structure, see Figure 1). We recorded the EEG brainwaves during both the pre- and post-learning face perception task. To maintain attention, participants pressed a button on a keyboard when an object was presented on the monitor (target hit rates > 0.99). In the explicit rating task, participants rated each of the 80 faces with a mouse on attractiveness, confidence, perceived competence, and perceived warmness (1 to 11).

The learning phase included a minimal group formation task, an associative learning task, and a social learning task with EEG recording. In the minimal group formation task, participants were randomly assigned to one of two groups (Green or White) and were told that the group assignment was based on the similarity of their personal preferences with the others^29^. Participants then completed an associative learning task, in which they pressed a button as soon as possible when their name was paired with their assigned group labels, among other names and the other group label pairs. This task served to strengthen the learned associations between their names and their assigned group labels. Participants indeed showed higher in-group identification and favoritism (*p*s < .002, Data S6). During the social learning task, participants were presented with the face again, together with the attractiveness rating feedback from either in-group or out-group members (i.e., Affiliation), which was either HIGHER, LOWER, or CONSISTENT (i.e., Feedback) than/with their initial ratings, resulting in a 2 (Affiliation) by 3 (Feedback) within-subject design. Assignments of experimental faces to each of the six conditions were counterbalanced across participants, with 10 additional faces as no-learning control faces. The post-learning phase was the same as the pre-learning phase, except that participants completed a cued recall task on their memories of the faces and the feedback before the perception and the rating tasks. Participants then provided their demographic information and answered group identification questions. Details of the minimal group manipulation, cued recall task, and social learning task are provided in Supplemental Methods.

Seven days later, participants visited the lab for the delayed phase. EEGs were not recorded in this phase. Analyses of the learning task and the cued recall tasks are beyond the scope of the current experiment and are not reported here.

### EEG Acquisition and Preprocessing

Continuous EEGs were recorded with an eego amplifier and a 64-channel gel-based waveguard cap based on an extended 10–20 layout (ANT Neuro, Enschede, and Netherlands). The online sampling rate was 500 Hz. The online reference electrode was CPz, and the ground electrode was AFz. The horizontal electrooculogram (EOG) was recorded from an electrode placed 1.5 cm to the left external canthus. The impedance of all electrodes was maintained below 20 kΩ during the recording.

For EEG data from the face perception task, we excluded trials in which participants accidentally pressed the button to faces. EEGs were processed offline using custom scripts, the EEGLAB toolbox^72^, and the ERPLAB toolbox^73^ implemented in MATLAB (MathWorks Inc, Natick, MA, USA). Raw EEG signals were first downsampled to 250 Hz and bandpass-filtered in the frequency range of 0.05–30 Hz using the FIR filter implemented in EEGLab. We removed 50 Hz line noise by applying the CleanLine algorithm^74^. EOG, M1, and M2 electrodes were removed from the EEG data before further processing. Bad channels were visually detected, removed, and then interpolated. To facilitate the independent component analysis (ICA) by including more datapoints (i.e., longer epochs), the EEG data were segmented into [-1000 to 2000 ms] epochs relative to the onset of the face and were then high-pass filtered with a cutoff frequency of 1 Hz^75^ before being subjected to ICA. After the ICA, artifacts caused by eye movements and muscle activity were identified and corrected using visual inspection and the ICLabel plugin^76^ implemented in EEGLAB. In addition, artifacts were automatically identified using the threshold of +/− 100 μV. Note that we preregistered a threshold of +/− 75 μV to exclude EEG artifacts. However, adopting this stricter threshold resulted in more excluded trials, thus reducing statistical power (Table S1). The results remained similar when using the preregistered preprocessing threshold (Data S7 and Table S4). Trials with artifacts or with incorrect responses (i.e., false alarms) were excluded from further analysis. On average, 475.50 (*S.D.* = 34.13) and 452.99 (*S.D.* = 44.01) trials were included for pre- and post-learning face perception EEG analyses, respectively.

### Event-Related Potential (ERP) Analysis

For ERP quantifications, continuous EEGs were segmented into [-200 to 1000 ms] epochs and were averaged for ERPs using the −200-0 pre-stimulus as baselines (preregistered). We decided only to include a shorter window because face and attractiveness perception is usually rapid and automatic^77^. We preregistered our intention to analyze the face-processing component N170^33^ at pre-defined bilateral occipitotemporal sites (left: T7, TP7, P7, PO7; right: T8, TP8, P8, PO8), and the evaluation-related component LPC^32^ at the pre-defined central-parietal (CPz, CP1/2, Pz, P1/2) and frontal-central (Fz, FCz, F1/2, FC1/2) sites. We measured 1) the mean amplitude for the time window of interest for each ERP component and 2) the adaptive mean by first finding the peak within the corresponding time window of interest and then calculating the mean around the peak^78^. The adaptive mean was based on the mean amplitudes of a 50 ms time window for the N170 and of a 100 ms time window for the LPC. We conducted statistical analyses on the changes of N170 at left and right occipital-temporal sites and the changes of LPC at central-parietal and frontal-central sites (post-minus pre-learning N170/LPC mean amplitude).

### Multivariate Representational Similarity Analysis (RSA)

We calculated the neural representation similarity between the experimental face and prototype attractive faces, i.e., the experimental-prototype face similarity (EPS) as a neural index of attractiveness. EEG data were downsampled to 100 Hz to facilitate multivariate similarity analyses. To reduce the effect of ERP on the multivariate RSA, we applied the z-transformation to the EEG activities: All individual EEG trials were normalized by subtracting the mean and were then divided by the standard deviation of ERP activities at each time point within each participant^79^. Next, the −200 to 1000 ms EEG epochs were continuously segmented into overlapping windows of 200 ms with 10 ms increments. By Spearman Correlation, we calculated the neural pattern similarity between individual time windows of every two trials (experimental face and prototype face) across all 61 channels. To control the temporal proximity effect (i.e., higher similarities would be expected for adjacent trials), we only analyzed the trials with more than four trials apart. The similarity of each face was averaged across all the correlation coefficients between all the trials of this face and all prototypical attractive faces. We conducted the cluster-based non-parametric permutation test by shuffling the subject label and constructing a null distribution 5,000 times with the default functions implemented in FieldTrip^80^.

### Statistics and reproducibility

For behavioral results, we conducted all repeated measures analysis of variance (ANOVA) using afex packages implemented in R, and post-hoc analysis using R package emmeans with bonferroni methods for multiple comparison. To provide more information out of null results, we further conducted bayesian analysis using BayesFactor implemented in R. For the statistical analysis of the RSA, we conducted nonparametric cluster-based permutation test with following parameters: 5000 permutations, two-tailed for *t*-test, cluster thereshold of *p* < 0.05, and a final threshold of *p* < 0.05 using fieldtrip toolbox^80^.

## Data and code availability

The pre-registration, data, and analysis scripts are publicly accessible at OSF and can be accessed at https://osf.io/5e7kr/?view_only=cf903bc29f8543a19272046a45a8349c https://osf.io/cg6rn. Deviations from pre-registration and corresponding reasoning can be found in Table S1.

## Acknowledgments

We thank Hui Xie for providing the stimuli dataset, Winny W.Y. Yue for her assistance in data collection, and Ruoying Zheng for her comments on the early draft.

## Author contributions

**D.C.:** Conceptualization, Investigation, Formal Analysis, Data Curation, Software, Methodology, Writing - Original Draft, Writing - Review & Editing, Visualization; **Z.Y.:** Conceptualization, Validation, Writing - Review & Editing; **J.L.:** Methodology, Writing – Review & Editing; **H.W.:** Conceptualization, Writing – Review & Editing; **X.H.:** Conceptualization, Writing - Original Draft, Writing - Review & Editing, Supervision, Project Administration, Funding Acquisition.

## Fundings

The research was supported by the Ministry of Science and Technology of China STI2030-Major Projects (No. 2022ZD0214100), National Natural Science Foundation of China (No. 32171056), General Research Fund (No. 17614922) of Hong Kong Research Grants Council, and the Key Realm R&D Program of Guangzhou (No. 20200703005) to X. H.

## Competing interests

The authors declare no competing financial or non-financial interests.

## Footnotes

1 We conducted the sensianalysis with G*Power ^63^. As we focused on the main effect of feedback, we conducted a sensitivity analysis with number of groups = 1, measurements = 3, and sample size = 42. The lowest power was set as 0.80, while the highest power was set as 0.95.

## Notes

**Conflict of Interest Statement:** The authors declare no conflict of interest.

### Competing Interest Statement

The authors have declared no competing interest.

### Summary of Updates

Title to be clearer; Introduction to provide more theoretical backgrounds; Results to be clearer; Discussion to be more accurate; Methods to provide more details.

