## Supplementary material for "Social Conformity Updates the Neural Representation of Facial Attractiveness": Fig S1

### Supplementary Materials

Danni Chen[1], Ziqing Yao[1], Jing Liu[2], Haiyan Wu[3] and Xiaoqing Hu[1,4*]

1, Department of Psychology, The State Key Laboratory of Brain and Cognitive Sciences, The University of Hong Kong, Hong Kong SAR, China

2, Department of Applied Social Sciences, The Hong Kong Polytechnic University, Hong Kong SAR, China

3, Centre for Cognitive and Brain Sciences and Department of Psychology, University of Macau, Macau SAR, China

4, HKU-Shenzhen Institute of Research and Innovation, Shenzhen, China

### Supplemental Methods

**Cued-Recall Task.** Participants completed the cued-recall task in the post-learning and delayed session, during which participants reported whether they have viewed the face stimuli in the learning task, and the group ratings. Each trial started with a blank display (0.5 - 0.9 s), followed by face stimuli (1 s). Then participants were asked to indicate whether they have seen the face stimuli in the previous learning session by pressing 'D' on the keyboard, otherwise pressing 'K'. Next, participants shall use the mouse click on the group ratings presented in the Learning Task (1 – 11).

**Minimal group formation task with associative training.** To manipulate group affiliations in a lab setting, we employed the classic minimal group formation task (Otten, 2016; Tajfel et al., 1971). In this minimal group formation task, participants answered three questions regarding their personal preferences:1) cats or dogs, 2) hamburgers or pizza, and 3) big cities or small towns (Goldenberg et al., 2020), and were randomly assigned into one of two groups: namely the "Green" group or the "White" group. They were told that the group assignment was based on the similarity of their personal preferences with the others. Participants were told that the assignment was merely based on the similarity of their preference between themselves and members of that specific group. To reinforce a sense of belongingness, participants 1) were instructed to wear a bracelet (i.e., a green bracelet or white bracelet) that indicated their group affiliation and 2) to complete an associative training task.

In the associative training task (revised from Yuan et al., 2019), participants were asked to press a button when their given name and corresponding group label appeared simultaneously on the screen.  First, participants spent a minute learning the group affiliations of six new names, with half belonging to the green group and the other half to the white group. Following this, a name (participants' own name, two names from the same group, and three names from the other group) and its related group name were displayed on the screen (e.g., Danni + Green). The other names were included as fillers. The associative training task consisted of four blocks, each containing five repetitions. Every repetition included one name from the green group, one from the white group, and the participant's own name paired with both the green and white groups. Participants had to press a button to determine if the name and group matched within 2000 ms

("D" for match, "K" for mismatch), after which they would receive feedback on their accuracy within 1000 ms. The association task aimed to enhance further the associative strength between participants' identities and their group affiliations.

Our group manipulation was successful, evidenced by the fact that 1) all participants correctly reported their group affiliation by the end of the first lab visit, 2) participants reported significantly higher identification, perceived positive characteristics, and perceived level of closeness to the ingroup members than outgroup members (one-tailed paired t-test, $p$s < .001, $d$s > 0.36, $BF_{10}$ > 17.44; Details see Data S6).

**Learning task.** During the learning task, participants learned the face attractiveness ratings as feedback from others. The learning included 300 trials, presented in 5 blocks, with each block containing 60 images from the six conditions (ingroup-higher, ingroup-lower, ingroup-consistent, outgroup-higher, outgroup-lower, outgroup-consistent). Each trial started with a blank screen (500 – 900 ms), followed by a fixation cross (500 ms). The face photo was then presented in the center of the screen for 1500 ms, together with participants' initial attractiveness rating from the pre-learning session, indicated by the same blue triangle cursor on the attractiveness rating scale (1500 ms). Following a blank screen (1000 ms), the same face photo and participants' initial attractiveness rating appeared again, with an additional rating of the averaged attractiveness in either a green (indicating green group ratings) or a white circle (indicating white group ratings, 3000 ms). As participants had been assigned to either green or the white group at the minimal group manipulation stage, the color of the circle thus represented in- or out-group feedback. Upon learning about others' ratings, participants rated the attractiveness again (3000 ms maximum) using the mouse. To encourage participants to pay attention to the group rating, they were told that their memory for the group rating would be tested later.

The group attractiveness ratings were pre-programmed for each participant. Ratings were either consistent, higher, or lower than participants' initial ratings from the pre-learning session. One-third of the feedback was consistent with participants' baseline attractiveness rating (CONSISTENT condition). For the remaining two-thirds of the feedback, half of the feedback was higher than the participants' first attractiveness rating (HIGHER condition), while the other half of the feedback was lower (LOWER condition). In the higher or lower conditions, the group ratings would be 1, 2, or 3 points above or below the participants' initial ratings, respectively. To increase the authenticity, the chance of 3 points difference was less likely than that of a 1 or 2 points difference.

### Supplemental Data

**Data S1. Delayed Updates of Attractiveness Rating from pre-learning to delayed sessions**. We also performed 2 by 3 repeated measure ANOVA was conducted on the update of mean-corrected attractiveness rating from pre-learning to the delayed session. No significant main effect of affiliation ($F$ (1, 45) = 0.30, $p$ = .587, $\eta^2$ = 0.002, $BF_{10}$ = 0.16) and feedback ($F$ (1.95, 87.61) = 0.84, $p$ = .430, η2 = 0.007, $BF_{10}$ = 0.09) was found. No significant interaction effect was found either ($F$ (1.94, 87.33) = 0.43, $p$ = .645, $\eta^2$ = 0.003, $BF_{10}$ = 0.16). Next, we conducted two repeated measure ANOVAs for ingroup and outgroup conditions separately. Again, we did not observe any significant main effect of feedback for both ingroup ($F$ (1.88, 84.55) = 0.52, $p$ = .586, $\eta^2$ = 0.007) and outgroup ($F$ (1.95, 87.63) = 0.82, $p$ = .440, $\eta^2$ = 0.012) conditions.

**Data S2. Immdiate Updates of Attractiveness Rating from pre-learning to post-learning sessions matching baseline ratings.** Taking the regression to mean effect into consideration, we further replicate the analyses with selected subsets of facial stimuli for which the participant's baseline attractiveness ratings were matched across the higher, lower and consistent conditions for the ingroup conditions and the outgroup condition separately. The results remained the same: We observed significant main effect of feedback, $F$ (1.87, 88.04) = 7.69, $p$ = .001, $\eta^2$ = .056. No significant main effect of affiliation, $F$ (1, 47) = 0.60, $p$ = .442, $\eta^2$ = .002, and interaction effect were found, $F$ (1.95, 91.56) = 0.12, $p$ = .883, $\eta^2 < .001$.

**Data S3. Effect of feedback and affiliation on the EPS in the pre-and post-learning phases.** We first examined whether the EPS differed across different feedback (higher, lower vs. consistent) in the pre-learning baseline phase, for in- and out-group conditions respectively, which showed no significant difference in the EPS across different feedback conditions for both in- and out-group conditions (in-group: $p_{\text{cluster}}$s > .114;  out-group: $p_{\text{cluster}}$s > .587). We further examined whether there were differences between in- and out-group through an affiliation (in- vs. out-group) by feedback repeated measures ANOVA on the EPS, which did not reveal a significant difference ($p_{\text{cluster}}$s > .613).

Next, we examined whether the EPS differed across different feedback in the post-learning baseline phase, which revealed no significant effect in EPS of either in- or out-group influence in

EPS (in-group: $p_{cluster}$ = .069; out-group: $p_{cluster}$s > .466). Similarly, the affiliation by feedback repeated measures ANOVA no significant difference between in- and out-group conditions on EPS was observed ($p_{cluster}$s > .786).

**Data S4. The difference in in-group favoritism between High- and Low-TLQ groups**. To eliminate the possibility that the perceived tightness-looseness might influence the in-group favoritism and further influence the level of conformity, we conducted independent sample t-tests and found no significant difference between the high- and low-TLQ group in the in-group identification, perceived positive characteristic, and perceived closeness (*p*s > .526, *d*s < 0.19); or in-group minus out-group identification, perceived positive characteristic, and perceived closeness (*p*s > .144, *d*s < 0. 43).

**Data S5. Prototypical faces were more attractive than the target faces.** These faces were selected based on ratings from an independent sample of 18 participants. The pilot results justified that the prototypical faces were more attractive than the target faces (scale from 1 to 7, prototype faces, *mean* = 5.84, *S.D.* = 0.66; target faces, *mean* = 3.77, *S.D.* = 0.69; $t$ (17) = 11.91, $p$ < .001, $d$ = 3.09, $BF_{10} = 9.22 \times 10^{6}$). Moreover, the attractiveness rating of the prototypical faces was significantly higher than the middle point, $t$ (17) = 12.12, $p$ < .001, $d$ = 2.86, $BF_{10} = 1.19 \times 10^{7}$, while the attractiveness rating between experimental faces was not different from middle point (4), $t$ (17) = 1.40, $p$ = .179, $d$ = 0.33, $BF_{10}$ = 0.56.

**Data S6. Minimal-Group Manipulation Check.** To check whether our minimal group manipulation worked, we performed a one-tailed t-test, which indeed showed that higher identification (in-group, *mean* = 4.81, *S.E.* = 0.11; out-group, *mean* = 3.27, *S.E.* = 0.15; $t$ (47) = 7.69, $p$ < .001, $d$ = 1.71, $BF_{10} = 3.02 \text{ x } 10^{7}$), perceived positive characteristics (in-group, *mean* = 4.86, *S.E.* = 0.13; out-group, *mean* = 4.55, *S.E.* = 0.12; $t$ (47) = 3.04, $p$ = .002, $d$ = 0.36, $BF_{10}$ = 17.44), and perceived level of closeness (in-group, *M* = 4.79, *S.E.* = 0.18; out-group, *mean* = 3.67, *S.E.* = 0.20; *V* = 512, $p$ < .001, $d$ = 0.84, $BF_{10}$ = 6460.06) to the ingroup members than the outgroup members. The results justify the effectiveness of our group membership manipulation.

**Data S7. Pre-registered Exploratory ERP Analysis with a threshold of 75 μV.** We conducted a series of 2 by 3 repeated measures ANOVA on the update of N170 and LPC amplitude for each

ROI separately. For N170 (adaptive mean) collapsed across Right Occipital-temporal site, we found in-group condition would elicit higher N170 update across different feedback conditions than out-group condition, $F$ (1, 40) = 4.30, $p$ = .045, $\eta^2$ = .010. No other significant main effects of affiliation, and feedback, and interactive effects were found (Table S4).

**Data S8. Perceived warmness from pre-learning to post-learning sessions**. A 2 by 3 repeated measure ANOVA on the update of mean-corrected warmness rating from the pre- to post-learning sessions (Figure S5A) revealed a marginally significant main effect of feedback ($F$ (1.99, 93.39) = 3.05, $p$ = .053, $\eta^2$ = 0.020, $BF_{10}$ = 0.51). Post-hoc analysis with Bonferroni correction revealed that the update of mean-corrected warmness rating is marginally significantly higher in the higher condition than in the lower condition, $t$ (47) = 2.45, $p$ = .054, $d$ = 0.33. No significant difference between consistent condition and higher / lower conditions, $p$s > .284, $d$s < .254.

**Data S9. Perceived warmness from pre-learning to delayed sessions**. We conducted the same 2 by 3 repeated measure ANOVA on the update of mean-corrected warmness rating from pre-learning to delayed sessions (Figure S5B). No significant main effect of affiliation ($F$ (1, 45) = 0.09, $p$ = .761, $\eta^2$ < 0.001, $BF_{10}$ = 0.13) and feedback ($F$ (1.99, 89.40) = 0.03, $p$ = .966, $\eta^2$ < 0.001, $BF_{10}$ = 0.04) was found. No significant interaction effect was found either ($F$ (2.00, 89.94) = 0.61, $p$ = .545, $\eta^2$ = 0.006, $BF_{10}$ = 0.25).

**Data S10. Perceived competence from pre-learning to post-learning sessions**. A 2 by 3 repeated measure ANOVA on the update of mean-corrected competence rating from the pre- to the post-learning (Figure S5C) did not find any significant main effect of affiliation, $F$ (1, 47) = 0.46, $p$ = .500, $\eta^2$ = 0.003, $BF_{10}$ = 0.19, and of feedback, $F$ (1.96, 92.17) = 1.87, $p$ = .160, $\eta^2$ = 0.012, $BF_{10}$ = 0.19. The significant interaction effect was not observed either, $F$ (1.85, 86.83) = 2.52, $p$ = .090, $\eta^2$ = 0.017, $BF_{10}$ = 0.94.

**Data S11. Perceived competence from pre-learning to delayed sessions**. Similar 2 by 3 repeated measure ANOVA on the update of mean-corrected competence rating from the pre- to the delayed session did not find any significant main effect of affiliation, $F$ (1, 45) = 1.03, $p$

= .315, $\eta^2$ = .005, $BF_{10}$ = 0.24, and of feedback, $F$ (2.00, 89.84) = 1.56, $p$ = .215, $\eta^2$ = .013, $BF_{10}$ = 0.20. No significant interaction effect was found either, $F$ (1.78, 79.92) = 1.29, $p$ = .278, $\eta^2$ = .010, $BF_{10}$ = 0.39 (Figure S5D).

**Data S12. Correlations between the update of the attractiveness rating and perceived personality ratings.** To explore whether the update of attractiveness rating driven by social influence could generalize to the perceived personality ratings, we conducted Pearson correlation analyses between the update of the attractiveness rating and the update of perceived warmness and competence ratings. In the immediate test, the update of the mean-corrected attractiveness rating was positively correlated with the update of the mean-corrected warmness rating across group affiliations: $r$ = .17, $p$ = .017 (Figure S6). It was driven by the in-group condition, $r$ = .22, $p$ = .033, rather than the out-group condition: $r$ = .13, $p$ = .193. No significant correlations between the update of competence and attractiveness rating were found: $r$s < .10, $p$s > .215. These findings indicated that the social influence on attractiveness perception generalized to perceived warmness but not competence, an effect particularly stronger in the in-group condition.

**Data S13. Correlation between the update of the attractiveness rating and perceived personality ratings at the delayed session.** In the delayed test, no significant correlation between the update of mean-corrected attractiveness rating and warmness ratings was found, all, $r$ = .06, $p$ = .44, in-group, $r$ = .03, $p$ = .80, out-group, $r$ = .09, $p$ = .40. No significant correlation between the update of competence and attractiveness rating was found, all, $r$ = .12, $p$ = .10, in-group, $r$ = .10, $p$ = .33, out-group, $r$ = .15, $p$ = .17.

**Data S14. Gender differences.** Some prior research suggests that female participants are more sensitive to social influence (Eagly, 1983; but see Wijenayake et al., 2020) and show stronger ingroup bias than their male counterparts (Herlitz & Lovén, 2013; Rudman & Goodwin, 2004). Based on these findings, we examined whether the social conformity effect in explicit ratings differed between different genders.

In the immediate test, we conducted a 2 by 3 by 2 mixed measure ANOVA, with affiliation and feedback as within-group factors and gender as the between-group factor. We found a significant main effect of feedback, $F$ (1.90, 87.60) = 6.47, $p$ = .003, $\eta^2$ = .053, and a significant three-way interaction effect, $F$ (1.96, 89.98) = 3.66, $p$ = .031, $\eta^2$ = .026. To unwrap the three-way interaction effect, we conducted two 2 by 3 repeated measure ANOVA separately for female and male participants. For female participants, we found a significant main effect of feedback, $F$ (1.86, 66.83) = 8.47, $p$ < .001, $\eta^2$ = .091. Post-hoc analysis revealed that the update of the mean-corrected attractiveness rating in the lower condition was significantly lower than of the higher condition, $t$ (36) = 3.63, $p$ = .002, $d$ = 0.73, and than of the consistent condition, $t$ (36) = 2.65, $p$ = .031, $d$ = 0.48. No significant difference between the higher and consistent condition was found, $t$ (36) = 1.79, $p$ = .189, $d$ = 0.31. No significant main effect of affiliation and interaction effect was observed ($p$s > .167). However, for male participants, we did not find a significant main effect of feedback, $F$ (1.82, 18.22) = 2.44, $p$ = .119, $\eta^2$ = .075, and affiliation, $F$ (1, 10) = 0.61, $p$ = .452, $\eta^2$ = .015. No significant interaction effect was observed either, $F$ (1.86, 18.56) = 2.60, $p$ = .104, $\eta^2$ = .087. The same 2 by 2 by 3 mixed ANOVA analysis on the delayed test did not find any significant main effect and interaction effect, $p$s > .355.

**Data S15. Correlation between the fantasy scale and the update of attractiveness ratings.** In the immediate test, for the higher condition, the update of the mean-corrected attractiveness rating was positively correlated with the fantasy scale ($r$ = .21, $p$ = .04). We further conducted linear regression analyses taking fantasy scale, feedback (higher vs. lower), and their interaction as independent variables, and the update of mean-corrected attractiveness rating as the dependent variable. For fantasy scale, we found that fantasy scale was a significant predictor ($b$ = 0.14, $p$ = .025), and the interaction between feedback and fantasy scale was a marginally significant predictor ($b$ = -0.15, $p$ = .079). Post-hoc analysis revealed that the standardized regression coefficient for the higher condition is significantly higher than for the lower condition, $p$ = .079, $t$ (188) = 1.78.

**Data S16. Differences in the updates of attractiveness ratings among different discrepancy levels (group ratings minus baseline self-ratings).** To better understand how participants would change their attractiveness rating in response to the discrepancy levels (Huang et al., 2014; Kim et al., 2012), we examined how explicit behavioral conformity would vary as discrepancy levels changed. Considering the unbalanced numbers of items among different discrepancy levels, we conducted a condition-level linear mixed model, taking affiliation and discrepancies as fixed factors, the number of items within each discrepancy as covariates, and affiliations as the random slope. The model used was as follows:

Update of Attractiveness Rating ~ Affiliation * Discrepancy +
Number of Items + (1+Affiliation | SubjectID)

The results showed a significant main effect of discrepancy levels ($F$ (6, 515.42) = 5.02, $p < .001$; Figure S2). Consistent with the analyses reported in the main text, we did not find a significant main effect of affiliation or affiliation by discrepancy level interaction ($p$s > .445). Post-hoc analyses revealed that a discrepancy level of +3 significantly increased attractiveness ratings than discrepancy levels that were lower than 0 ($p_{tukey}$s < .030). Moreover, a discrepancy level of +2 also significantly increased attractiveness ratings than discrepancy levels of -1 ($p_{tukey}$s = .043). Thus, corroborating the main analyses, these results showed that participants changed their attractiveness ratings upon peers' attractiveness ratings, regardless of in- or out-group members.

**Supplemental Figures**

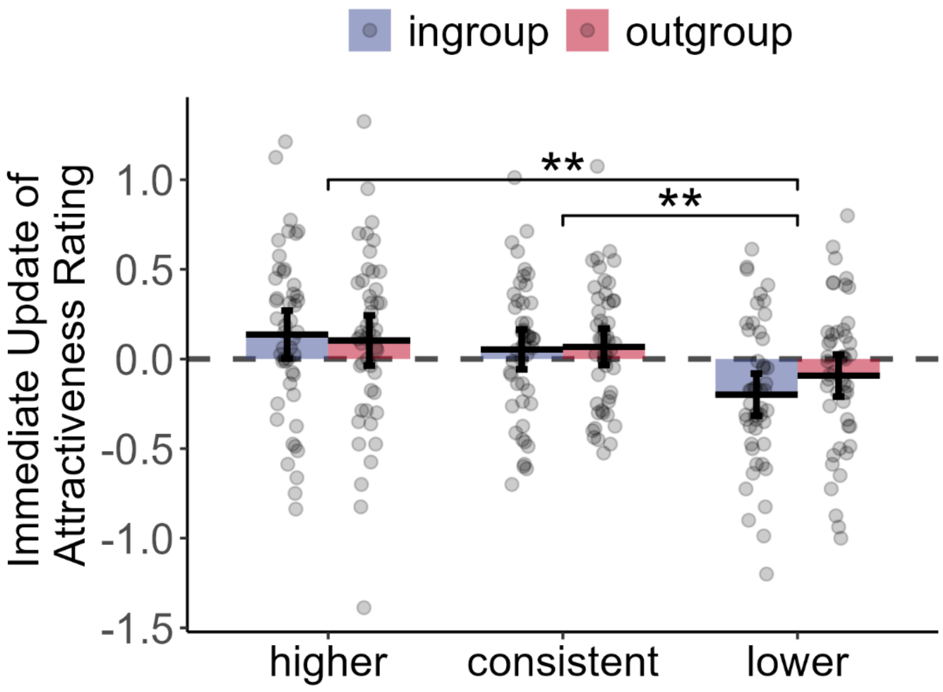

**Figure S1.** The effect of affiliation (in- and out-group) and feedback (higher, consistent and lower) on the updates of mean-corrected attractiveness ratings from pre- to post-learning phases. **: $p < .01$.

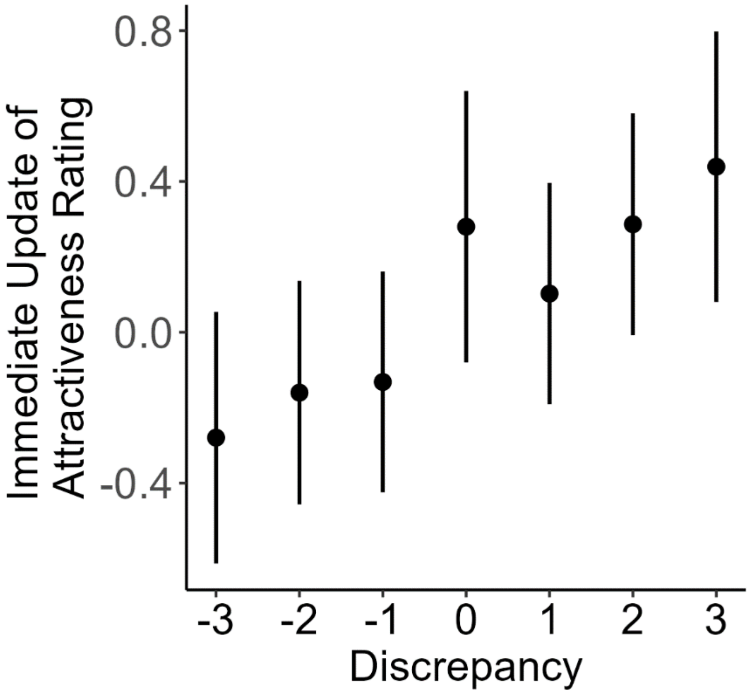

**Figure S2.** Differences in the immediate updates of attractiveness ratings from pre- to post-learning phase among different discrepancy levels (group ratings minus baseline self-ratings).

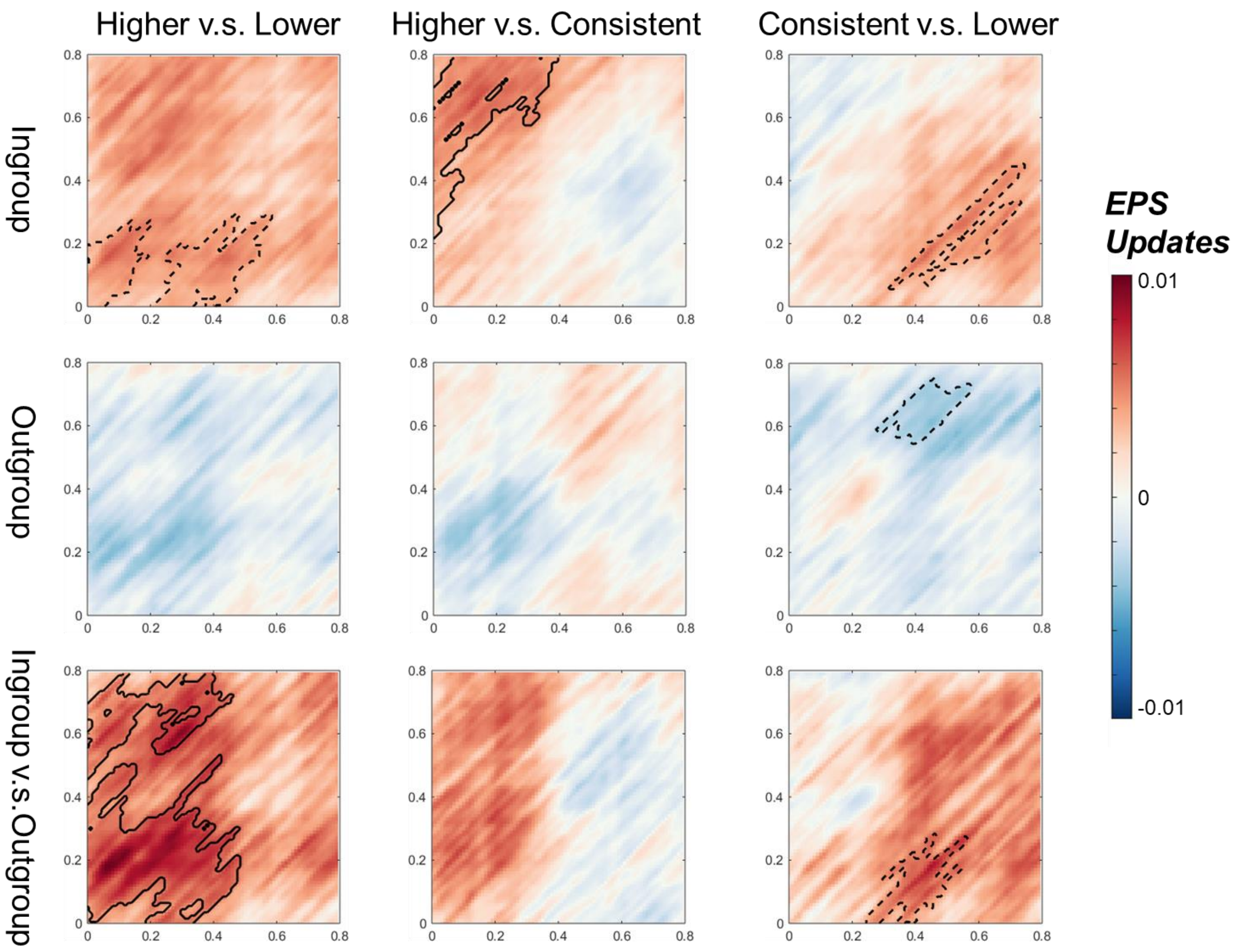

**Figure S3.** The RSA results for female participants (n = 37). We found significant clusters in the ingroup condition when comparing higher vs lower feedbacks. However, no significant positive clusters were observed. We further tested interaction effects by comparing ingroup vs outgroup. Results showed significant clusters of interaction effect when comparing higher vs consistent (*p*s < .041). The x-axis is the timescale of the experimental stimuli, and the Y axis is the timescale of the prototype stimuli. The dashed contour indicates marginally significant clusters ($p < .10$), while the solid contour indicates significant clusters ($p < .05$).

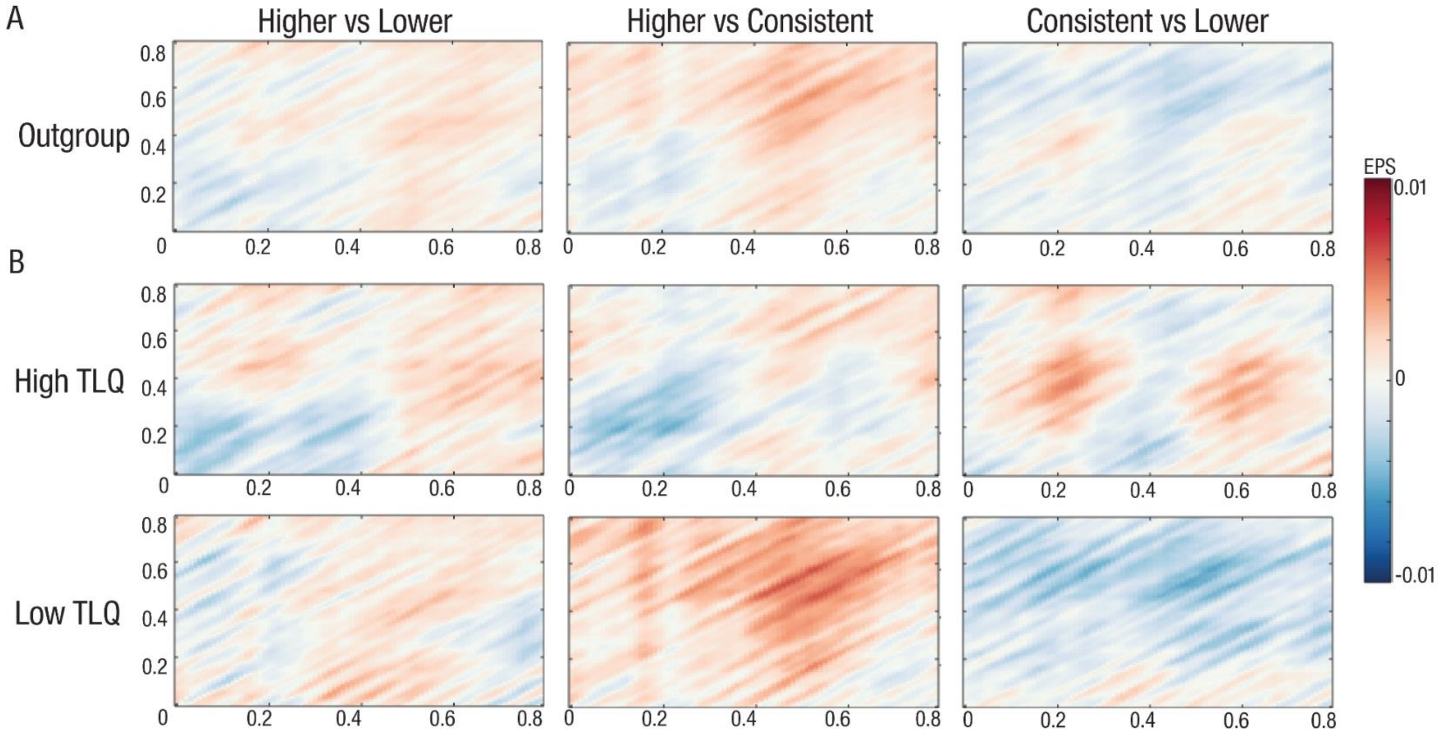

**Figure S4.** The RSA results in the outgroup conditions. A) The EPS update from the pre-learning to post-learning sessions in the outgroup conditions. No significant cluster was observed. B) The EPS update from the pre-learning to post-learning sessions in the High TLQ and Low TLQ sub-groups. No significant cluster was observed.

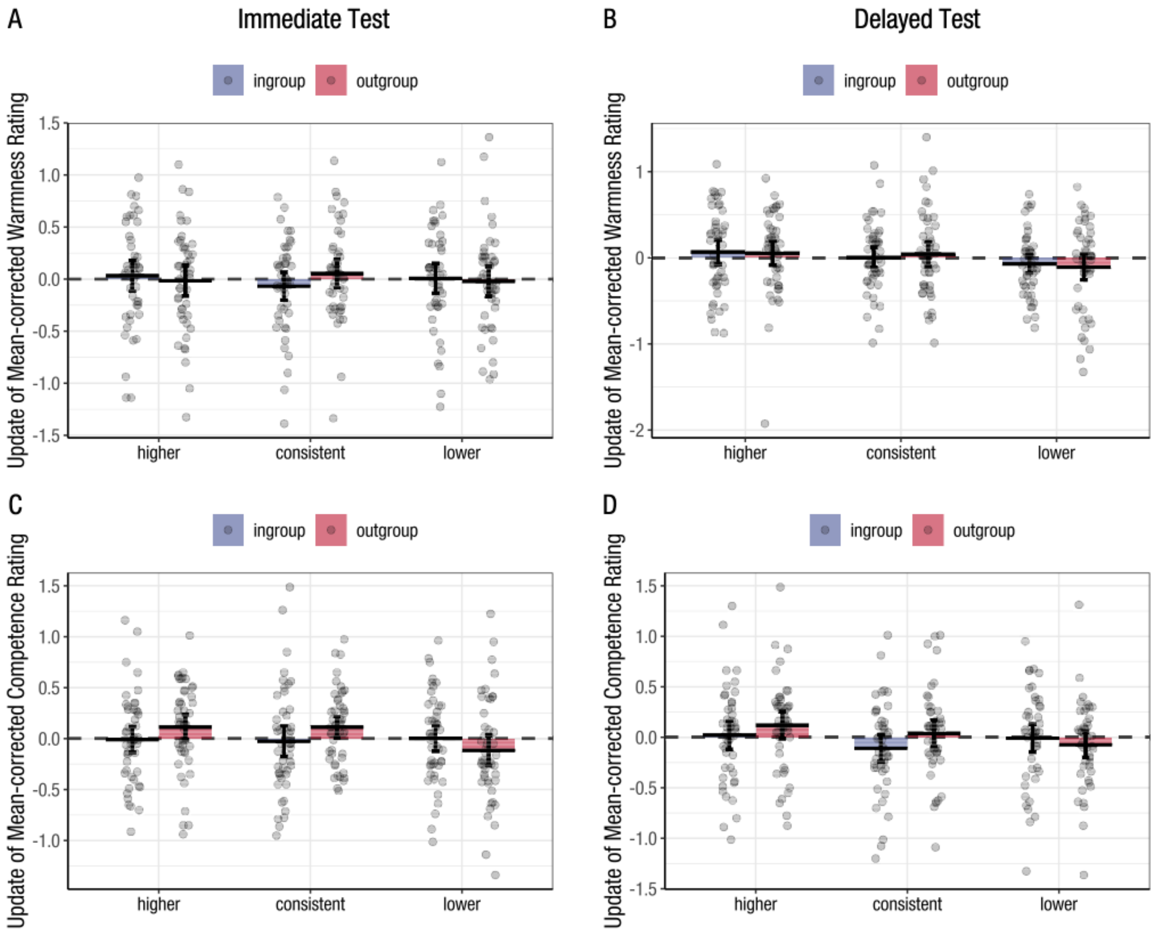

**Figure S5.** The update of mean-corrected warmness rating from the pre- to A) the post-learning session and to B) the delayed session. The update of mean-corrected competence rating from the pre- to C) the post-learning session and to D) the delayed session.

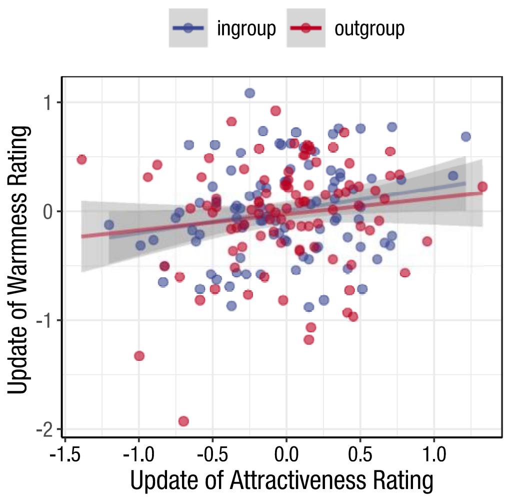

**Figure S6.** Correlation between Update of Attractiveness Rating and Update of Warmness Rating in the immediate test.

**Supplemental Tables**

**Table S1**

*Deviations from preregistration in the current EEG study.*

| Components in your preregistration | Were there deviations? What type? | If yes - describe details of deviation(s) | Rationale for deviation | How might the results be different if you had/had not deviated | Stage of decision for deviation |
|---|---|---|---|---|---|
| EEG Data Preprocessing | Yes. Minor. | We used CleanLine instead of the plugin implemented in the ERPLab. | CleanLine is a more updated Plugin for removing Line Noise and is recommended by the PREP pipeline (Bigdely-Shamlo et al., 2015). | NA | Before starting EEG data preprocessing. |
| | Yes. Minor. | We used the ICLabel Plugin instead of the ADJUST to help visually inspect the eye movement and muscle artifacts. | ICLabel is reported with higher classification accuracy than ADJUST (Pion-Tonachini et al., 2019). | NA | Before starting EEG data preprocessing. |

| | | | | | |
|---|---|---|---|---|---|
| | Yes. Intermediate. | We used 100 µV instead of 75 µV as the threshold to exclude trials. | Using 75 µV as the threshold, we would exclude more participants and more trials for each participant, which could reduce the power of the statistical analysis. With 100 µV as the threshold, we will retain 45 participants and 475.50 (S.D. = 34.13) and 452.99 (S.D. = 44.01) trials on average for pre- and post-learning sessions respectively. However, with 75 µV as the threshold, we will retain 41 participants and 436.32 (S.D. = 54.07) and 433.56 (S.D. = 56.34) trials on average for pre- and post-learning sessions respectively. | We have replicated the pre-registered ERP analysis for with the 75 µV (see Supplementary Material and Table S5). Most of the null effect of ERP remained the same. with the 75 µV, we found significant main effect of affiliation, feedback and interaction effect. However, the results were not consistent across regions of interest, and the effect sizes were small. It won't influence our conclusions. | During EEG data preprocessing but before any ERP-related statistical analysis. |

**Table S2.** Means, standard deviations, and correlations between attractiveness rating update and individual differences measurements with confidence intervals.

| Variable | *Mean* | *SD* | Higher | | Lower | |
|---|---|---|---|---|---|---|
| | | | Pre to Post | Pre to Delayed | Pre to Post | Pre to Delayed |
| ***IRI*** | | | | | | |
| FS | 3.46 | 0.72 | .21* | .07 | -.03 | .11 |
| | | | [.01, .40] | [-.14, .27] | [-.23, .17] | [-.10, .31] |
| EC | 3.47 | 0.58 | -.05 | .12 | .19 | .06 |
| | | | [-.25, .15] | [-.08, .32] | [-.01, .38] | [-.15, .26] |
| PT | 3.62 | 0.55 | .01 | .02 | .05 | -.04 |
| | | | [-.19, .21] | [-.18, .23] | [-.15, .25] | [-.25, .16] |
| PD | 2.97 | 0.55 | -.02 | .08 | .09 | .08 |
| | | | [-.21, .19] | [-.13, .28] | [-.11, .29] | [-.12, .28] |
| ***TLQ*** | 4.20 | 0.70 | .19 | -.03 | -.19 | -.04 |
| | | | [-.01, .37] | [-.23, .17] | [-.38, .01] | [-.24, .17] |
| ***SDR*** | | | | | | |
| SDE | 80.23 | 8.59 | -.11 | -.20 | -.00 | .00 |
| | | | [-.30, .09] | [-.39, .01] | [-.20, .20] | [-.20, .20] |
| IM | 87.94 | 9.20 | .01 | -.21* | -.05 | .03 |
| | | | [-.19, .21] | [-.40, -.01] | [-.25, .15] | [-.18, .23] |

| | | | | | | |
|---|---|---|---|---|---|---|
| ***SPIN*** | 40.02 | 10.27 | -.02 | -.04 | -.03 | .06 |
| | | | [-.22, .18] | [-.25, .16] | [-.22, .18] | [-.14, .26] |

*Note*. *M* and *SD* are used to represent mean and standard deviation, respectively. Values in square brackets indicate the 95% confidence interval for each correlation. IRI: Interpersonal reactivity index. FS: fantasy scale. PT: perspective taking. PD: personal distress. EC: empathic concern. TLQ: tightness-losseness questionnaire. SDR: Socially Desirable Responding. SDE: Self-deceptive enhancement. IM: Impression management. SPIN: Social Phobia Inventory. * indicates $p < .05$. ** indicates $p < .01$.

**Table S3**

*Results of 2 by 3 repeated measure ANOVAs on ERP amplitude updates from pre- to post-learning with a threshold of 100 µV*

| Component | ROI | Measurements | The main effect of Affiliation | | | | The main effect of Feedback | | | | Interaction effect | | | |
|---|---|---|---|---|---|---|---|---|---|---|---|---|---|---|
| | | | *F* | *p* | $\eta^2$ | $BF_{10}$ | *F* | *p* | $\eta^2$ | $BF_{10}$ | *F* | *p* | $\eta^2$ | $BF_{10}$ |
| N170 | ROT | M | 0.715 | .402 | 0.001 | 0.19 | 0.311 | .712 | 0.001 | 0.05 | 0.317 | .720 | 0.001 | 0.15 |
| | | A.M. | 3.867 | .056 | 0.008 | 1.68 | 0.023 | .976 | <0.001 | 0.04 | 0.993 | .370 | 0.003 | 0.25 |
| | LOT | M | 1.129 | .294 | 0.001 | 0.21 | 1.637 | .201 | 0.003 | 0.13 | 1.856 | .164 | 0.006 | 0.83 |
| | | A.M. | 0.567 | .455 | 0.001 | 0.16 | 1.647 | .199 | 0.004 | 0.15 | 2.562 | .088 | 0.009 | 1.74 |
| LPC | CP | M | 2.866 | .098 | 0.004 | 0.48 | 0.582 | .561 | 0.002 | 0.07 | 1.757 | .181 | 0.005 | 0.49 |
| | | A.M. | 2.532 | .119 | 0.005 | 0.48 | 0.79 | .445 | 0.003 | 0.08 | 2.703 | .073 | 0.008 | 0.87 |
| | FC | M | 0.409 | .526 | 0.001 | 0.16 | 1.494 | .231 | 0.004 | 0.17 | 0.538 | .582 | 0.001 | 0.18 |
| | | A.M. | 0.011 | .918 | <0.001 | 0.13 | 1.795 | .176 | 0.005 | 0.21 | 0.624 | .533 | 0.002 | 0.19 |

*Note*: CP: central-parietal sites (CPz, CP1/2, Pz, P1/2); OT: occipitotemporal sites (left: T7, TP7, P7, PO7; right: T8, TP8, P8, PO8); FC: frontocentral sites (Fz, FCz, F1/2, FC1/2). M: Mean; A.M.: Adaptive Mean.

**Table S4**

*2 by 3 repeated measure ANOVA on ERP Amplitude update results with a threshold of 75 μV*

| **Component** | **ROI** | **Measurements** | **The main effect of Affiliation** | | | | **The main effect of Feedback** | | | | **Interaction effect** | | | |
|---|---|---|---|---|---|---|---|---|---|---|---|---|---|---|
| | | | *F* | *p* | $\eta^2$ | $BF_{10}$ | *F* | *p* | $\eta^2$ | $BF_{10}$ | *F* | *p* | $\eta^2$ | $BF_{10}$ |
| N170 | LOT | M | 0.86 | .360 | <0.01 | 0.18 | 3.04 | 0.056 | 0.01 | 0.47 | 1.02 | .363 | <0.01 | 0.36 |
| | | A.M. | 1.10 | .300 | <0.01 | 0.19 | 2.96 | 0.063 | 0.01 | 0.61 | 2.07 | .137 | 0.01 | 0.92 |
| | ROT | Mean | 0.44 | .513 | <0.01 | 0.18 | 1.82 | 0.174 | 0.01 | 0.23 | 0.17 | .838 | <0.01 | 0.14 |
| | | A.M. | 4.30 | **.045*** | 0.01 | 3.05 | 0.35 | 0.699 | <0.01 | 0.06 | 0.81 | .447 | <0.01 | 0.22 |
| LPC | CP | M | 1.43 | .239 | <0.01 | 0.32 | 0.12 | 0.884 | <0.01 | 0.05 | 2.59 | .083 | 0.01 | 1.20 |
| | | A.M. | 1.42 | .241 | 0.01 | 0.33 | 0.24 | 0.773 | <0.01 | 0.05 | 3.10 | .051 | 0.02 | 1.79 |
| | FC | M | 0.14 | .707 | <0.01 | 0.15 | 2.64 | 0.082 | 0.01 | 0.43 | 0.41 | .658 | <0.01 | 0.18 |
| | | A.M. | 0.05 | .820 | <0.01 | 0.14 | 2.80 | 0.068 | 0.01 | 0.40 | 0.95 | .388 | <0.01 | 0.28 |

*Note*: CP: central-parietal sites (CPz, CP1/2, Pz, P1/2); OT: occipitotemporal sites (left: T7, TP7, P7, PO7; right: T8, TP8, P8, PO8); FC: frontocentral sites (Fz, FCz, F1/2, FC1/2). M: Mean; A.M.: Adaptive Mean. * indicates $p < .05$.
